# Sensitive identification of known and unknown protease activities by unsupervised linear motif deconvolution

**DOI:** 10.1101/2021.11.15.468703

**Authors:** Anuli C. Uzozie, Theodore G. Smith, Siyuan Chen, Philipp F. Lange

**Affiliations:** Department of Pathology, University of British Columbia, Vancouver, BC, Canada; Michael Cuccione Childhood Cancer Research Program, BC Children’s Hospital Research Institute, Vancouver, BC, Canada; Department of Molecular Oncology, BC Cancer, Vancouver, BC, Canada

**Keywords:** protease, N termini, substrate

## Abstract

The cleavage-site specificities for many proteases are not well-understood, restricting the utility of supervised classification methods. We present an algorithm and web interface to overcome this limitation through the unsupervised detection of overrepresented patterns in protein sequence data, providing insight into the mixture of protease activities contributing to a complex system.

Here, we apply the RObust LInear Motif Deconvolution (RoLiM) algorithm to confidently detect substrate cleavage patterns for SARS-CoV-2 Mpro protease in N terminome data of an infected human cell line. Using mass spectrometry-based peptide data from a case-control comparison of 341 primary urothelial bladder cancer cases and 110 controls, we identified distinct sequence motifs indicative of increased MMP activity in urine from cancer patients. Evaluation of N terminal peptides from patient plasma post-chemotherapy detected novel Granzyme B/Corin activity.

RoLiM will enhance unbiased investigation of peptide sequences to establish the composition of known and uncharacterized protease activities in biological systems.

## Introduction

Proteolysis affects most proteins in a cell. Protein function, localization or other properties are altered by proteolysis. Although advances in proteomics have enabled comprehensive identification of proteins in complex biological samples, the direct detection of protease activity remains challenging. This is due in part to (i) relatively low abundance of some proteolytic products, (ii) detection of a protease does not directly imply activity or co-localization with its substrate at the time of detection and (iii) simultaneous activity of multiple proteases. Hence, the detection of proteolytic activity is typically accomplished by observation of protease substrates pattern. This can be accomplished with experimental approaches involving termini enrichment^1^ or with activity based probes which directly recognize active proteases^2^.

Statistical algorithms are critical to derive biological insights from massive peptidomic datasets. Such algorithms should be able to address two main questions: (i) how accurately do peptide motifs identified in a data signify substrate cleavage or substrate cleavage pattern, and (ii) can detected sequence patterns be correctly linked to protease activity? Currently, there are two major strategies used for the indirect detection of proteolytic activity in large-scale peptide data generated by mass spectrometry. One approach is to predict putative protease cleavage sites based on observation of known cleavage events corresponding to a particular protease. Two popular algorithms that make use of this strategy are Proteasix and PROSPERous^3,4^. Here, supervised learning algorithms can be trained to recognize the sequence features characteristic of cleavage sites in the known substrates of a protease. Although this strategy has proven effective in certain contexts for some well characterized proteases^3,4^, it fails to account for the activity of less-understood proteases. Moreover, cleavage event data have been compiled over a substantial period of time via a shifting landscape of methodologies which vary drastically in both comprehensiveness and reliability. This strategy’s reliance on previous, curated data for the training of supervised learning algorithms makes it vulnerable to data that are either no longer current, relevant, or accurate.

The second approach relies on unsupervised motif detection for the analysis of existing sequence determinants of post-translational modifications (PTMs). These algorithms notably include Motif-X/Momo and GibbsCluster^5–7^. Momo supports the analysis of data sets containing multiple sequences with fixed central residues and these central residues are usually the target of a covalent modification such as phosphorylation^6,7^. Although these tools are promising resources for the unbiased analysis of PTM data centered around a modified residue, they are not suitable for the analysis of proteolytic data sets. Unlike other PTMs, proteolytic events are centered around a bond between two residues. Moreover, the amino acids found adjacent to proteolytic cleavage sites are highly variable and depend on the specific protease responsible for a cleavage event. Proteolytic data sets therefore lack the fixed central residue required by these tools, even if the sequence alignment is shifted in order to center the sequences around a residue position rather than the cleavage site.

Gibbs clustering, a popular Markov chain Monte Carlo algorithm used for clustering of sequence data sets^8^, is among the few algorithms that can align and cluster amino acid sequences without restriction to specified central residues. The GibbsCluster algorithm (GibbsCluster 2.0) relies on a non-deterministic algorithm (Gibbs sampling). This limits the reproducibility of results between runs and poses a major challenge to its use for analysing proteolytic input data, where the mixture of activities contributing to the composition of a sample is complex and the number of different protease activities (clusters) not known a priori.

In the present study, we extend a motif deconvolution algorithm we recently developed, RObust LInear Motif Deconvolution (RoLiM)^9^ to overcome the challenges associated with the unsupervised detection of protease activity in complex biological samples. We evaluate the performance of RoLiM in comparison to Proteasix and Gibbs clustering. In a re-analysis of protein N termini enriched from SARS-Cov2-infected human cell lines we demonstrate that RoLiM can identify new, uncharacterized protease activities. In re-analyses of urine collected from patients with urothelial bladder cancer, and extracellular fluids collected post chemotherapy from hematopoietic cancer patients we demonstrate that RoLiM can identify multiple cancer associated protease activities in complex extracellular fluids, and accurately assign identified protease patterns to known proteases or protease families.

## Results

We recently developed the Robust Linear Motif Deconvolution (RoLiM) algorithm for unbiased detection of statistically enriched amino acid patterns, and demonstrated that it exceeds existing solutions in robustness and versatility^9^ . We further demonstrated its application to the analysis of over 60 covalent protein modifications across 30 human tissues.

Here, we set out to explore RoLiM’s application to protein cleavage controlled by proteases.

### Identification of unknown proteolytic activities by unsupervised RoLiM analysis

We first tested the hypothesis that unsupervised analysis of protein cleavage sequence data enables the detection of unknown proteolytic activities missed by algorithms trained on existing knowledge. The MEROPS database^10,11^ is considered the most comprehensive repository of protease/peptidase information. It contains 92,216 substrate cleavage site information for 5267 peptidases^11^ in the form of substrate sequences with lengths of 8 amino acids.

Curated protease databases, in particular MEROPS, remain the backbone of many protease prediction algorithms. This presents a major limitation that centers on the inability of such algorithms to (i) detect enriched substrate patterns that are not categorized in the database, and (ii) match patterns to identify activities from proteases lacking curated cleavage site information. We benchmarked the performance of RoLiM with Proteasix which we believe is the most comparable tool for the prediction of protease activity in comprehensive peptide datasets.

We first evaluated the ability to detect the activity of an unknown protease in a complex system. We used the newly emerged SARS-CoV-2 virus as a model system in which the viral protease characteristics are known, but masked from the algorithms. Meyer and colleagues^12^ studied proteolysis induced by SARS-CoV-2 infection in cell lines using a mass spectrometry-based N termini enrichment method^13^. We tested the ability of unsupervised (RoLiM) and supervised (Proteasix) algorithms to predict protease activities from N termini quantified in A549-Ace2 human lung cell line pre- and post- SARS-Cov2 infection (Supplementary Table 1, Supplementary Fig. 1A, see Materials and Methods for details). We further sought to determine the capability of each algorithm to impartially identify substrate cleavage patterns that appear at defined timepoints after viral infection, and to possibly pinpoint likely proteases responsible for these proteolytic activities. Both approaches identified apoptotic signatures from caspases and granzymes 6-hr post infection (Figs. 1A and 1B).

**Figure 1.**
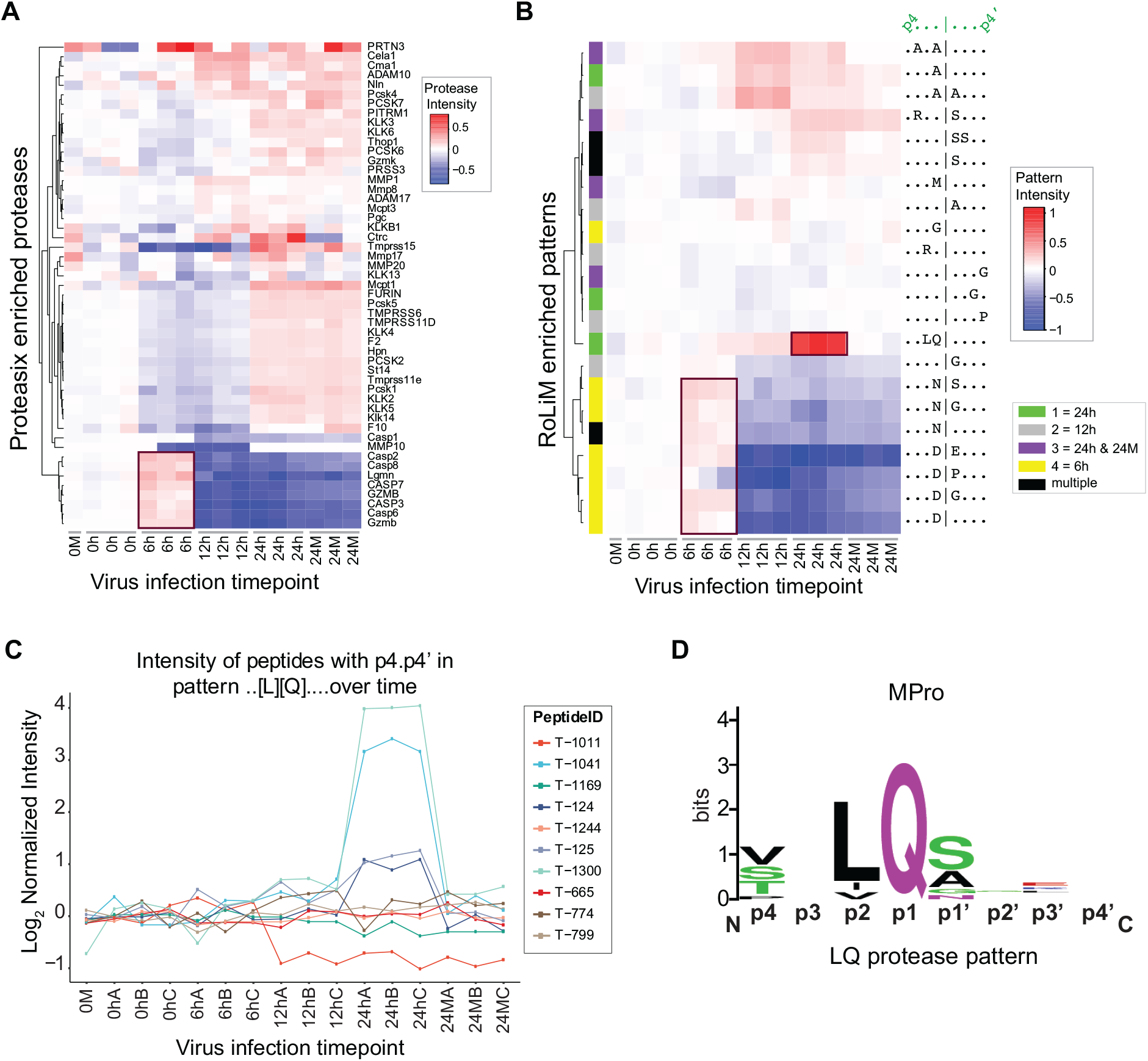
Unbiased detection of unknown sequence patterns in SARS-Cov2 infected human cell line. **A**. Heatmap on normalized peptide intensity ratios for protease activities predicted by Proteasix at different time points after SARS-Cov2 infection of human A549-Ace2 cell line. See Methods for details of the analysis. **B**. Heatmap on normalized peptide intensity ratios for enriched patterns detected by RoLiM at specific time points after viral infection. Row wise hierarchical clusters group patterns unique to one or more timepoint post-infection. **C**. Line plot shows the normalized intensity of peptides with the ..[L][Q]…. pattern in infected (6 h, 12 h, 24 h) and non-infected (0M, 24M) cells at all timepoints studied. **D**. Aligned sequence logo for SARS-CoV-2 MPro protease documented in MEROPS.

In addition, RoLiM found enriched patterns after 12 h and 24 h respectively (Fig. 1B and Supplementary Fig. 1B). At the 24 h timepoint, RoLiM detected a statistically enriched substrate motif (LQ|X) that could not be confidently assigned to any human protease in the MEROPS database. The intensity profile of the LQ|X pattern was clearly distinct from all other detected patterns (Fig. 1B). The intensity of individual peptides contributing to the LQ|X motif is shown in Fig. 1C. Meyer et al further validated the LQ|X pattern identified by RoLiM to be unique to the viral protease MPro (Fig. 1D). This analysis revealed the complete reliance on curated information in the MEROPS database by Proteasix as a key limitation. Proteasix also cannot identify cleavage events and peptide signatures for proteases with no known substrate cleavage information as well as proteases not associated with the organism(s) of interest. RoLiM, but not Proteasix, detected an activity that could be linked to infection by the novel SARS-CoV-2 virus. This confirms our hypothesis that unsupervised analysis of protease cleavage sequences enables the unbiased detection of unknown and unexpected cleavage patterns irrespective of organism specificity and prior knowledge.

### Implementation of proteolysis-specific functionality in RoLiM

We previously described the RoLiM algorithm in detail and evaluated sensitivity, false discovery rates, and robustness against changes in data composition^9^. Compared to existing algorithms for site-specific motif detection, particularly, Motif-X and Momo, RoLiM enabled a more robust and unbiased characterization of PTM modification sequence motifs, including phosphorylation, in large complex biological datasets. RoLiM further supports even-length sequence alignment centered around a (cleaved) peptide bond. To evaluate its ability to characterize protease substrate specificities we first sought to characterize the MEROPS human protease substrate data using RoLiM. MEROPS documents 884 known and putative human peptidases, including about 520 identified peptidases. Yet of these, we found that only 287 have at least one known physiological substrate and only 132 have more than 5 (Fig. 2A) clearly demonstrating our limited knowledge, even for a well-studied organism.

**Figure 2.**
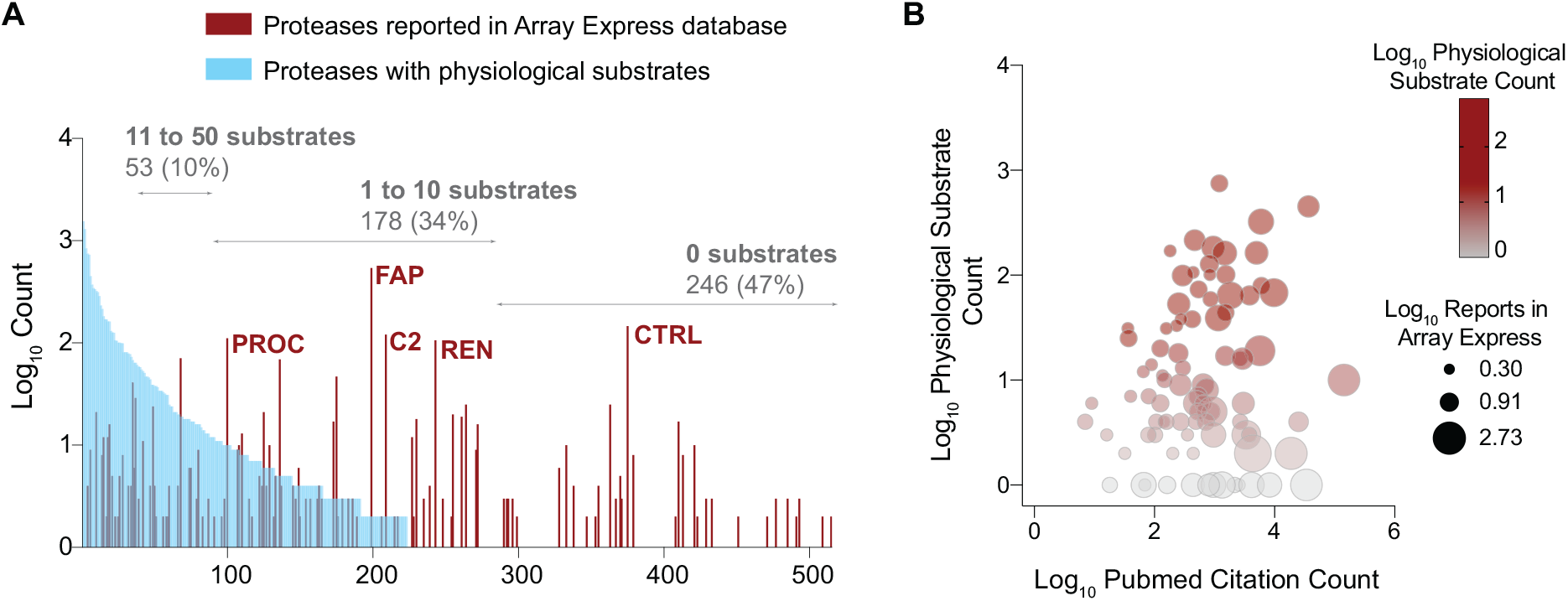
MEROPS human proteases. **A**. Barplots show the number and percentage of 520 curated MEROPS human proteases with known physiological substrates (blue bars) along with the number of times a protease has been identified and/or characterized in ArrayExpress, a publicly accessible repository for high-throughput proteomics and genomics data (brown bars). The five proteases with the highest counts in ArrayExpress are highlighted. **B**. Bubbleplot shows the poor relationship between proteases having known physiological substrates, the number of literature citations for these proteases in PubMed and the depth of functional studies reported in ArrayExpress.

We next reviewed two major public repositories - PubMed^14^ to retrieve the number of scientific literature on each human protease, and ArrayExpress^15^ to tally how often each protease has been reported in high-throughput functional genomics datasets (Supplementary Table 2). The results shown in Figs. 2A and 2B underscore the deficit in existing knowledge on human protease substrates, for both well-studied and poorly characterized proteases.

For further analysis, our input data consisted of sequences retrieved from the MEROPS database, filtered for “human active” protease substrates (indicating confirmed activity in humans), and synthetic and theoretical peptides were removed. Using the default settings for our algorithm (see Methods), we detected 261 patterns for 108 human proteases across all catalytic mechanisms (Supplementary Fig. 2A). We found that less than 20% of the 261 patterns were non-discriminatory while 70% of patterns differentiated between catalytic types and about 50% of the patterns was even able to discriminate different proteases of the same family (Supplementary Fig. 2b).

To enrich the unsupervised analysis of RoLiM with annotation based on existing knowledge - one of the key benefits of supervised algorithms, we decided to match patterns identified in the input data set to patterns associated with MEROPS proteases. To provide a comprehensive view of matched proteases and enable identification of unique pattern-protease matches as well as highly unspecific patterns or proteases we developed the protease substrate clustermap and heatmap (Supplementary Fig. 2C). RoLiM now provides users with a simple interface and processing steps for highly robust and sensitive unsupervised identification of proteolytic activities combined with intuitive annotation of known activities (Supplementary Figs. 1A and 2C).

### Improved extraction of distinct protease patterns from mixed protease activity data

We next assessed the ability of RoLiM and Gibbs clustering (GibbsCluster 2.0), an unsupervised motif discovery tool^16^, to correctly reject spurious patterns in a random sequence set that does not contain any statistically overrepresented patterns. The GibbsCluster algorithm takes a prespecified number of clusters and initializes the clustering workflow by randomly assigning each input sequence to one of those clusters. Clusters are represented by a stacked bar chart representing the Kullback Leibler Divergence (KLD) of each cluster and, cumulatively, the KLD of the system, as well as by a logo map corresponding to each pattern. To compare the performance of RoLiM and GibbCluster (v 2.0), we analyzed an *in silico* designed set of 1000 random 8 residue long sequences derived from the Swiss Prot human proteome (Supplementary Table 3, see Methods). Gibbs clustering selected an optimal result of 15 enriched patterns (average system KLD = 2.893) and respective logo maps, equal to the maximum number of clusters tested in this experiment (Supplementary Figs. 3 and 4). RoLiM on the other hand produced an empty result set using the same input data and similar settings confirming its robustness against random noise.

Following this, we compared the capability of RoLiM and Gibbs clustering to discern different patterns using a defined *in silico* mixture of actual protease substrate sequences. This comprised 964 sequences generated from 194 substrate sequences of caspase-6, 469 substrate sequences of cathepsin B, and 301 substrate sequences of matrix metallopeptidase 9 (MMP-9) derived from MEROPS. To simulate background noise, 987 theoretical peptides built from random alignment of 8 amino acids were also included (Supplementary Table 4). As shown in Fig. 3A, the caspase-6 subset is characterized by enriched acidic residues, particularly in the p4, p3 and p1 positions, along with valine in the p4 position. The cathepsin B substrate set is characterized by enrichment of small and hydrophobic residues with strong overrepresentation of glycine in positions p1 and P3’ respectively (Fig. 3B). Substrates for MMP-9 also displayed small and hydrophobic residues in several positions (Fig. 3C). Based on sequence composition, we anticipated that caspase-6 and cathepsin B substrates would be relatively easy to separate. While same would hold for cathepsin B and MMP-9, similarities in the amino acid composition of their substrates would prove challenging to a clustering-based approach when the subsets are combined. Logo maps corresponding to the mixed substrate data without and with background noise are presented in Figs. 3D and 3E respectively. These underscore the profound similarity of MMP-9 and cathepsin B substrates. Gibbs clustering was performed on the final data set with background noise using settings similar to the defaults in our algorithm (See Methods). Gibbs Clustering selected a solution of two clusters, with an average system KLD of 3.432 (Fig. 3F). Even if forced to separate into three clusters based on prior knowledge of three mixed activities, Gibbs clustering failed to resolve MMP-9 and cathepsin B substrates into separate clusters (Figs. 3G-I). RoLiM detected patterns in the mixed sample that mapped effectively to known MEROPS patterns from the three proteases used to compose the test dataset. As shown in Fig. 3J, distinct pattern matches were observed for cathepsin B as well as multiple caspases and matrix metallopeptidases including caspase-6 and MMP-9. Sequence logos for enriched patterns are depicted in Figs. 3K to N. The same patterns matching these proteases also matched other proteases from similar MEROPS families. This is a consequence of the fact that proteases from the same family often exhibit extremely identical, if not similar cleavage site specificity. These results demonstrate the efficacy of the pattern detection performed by our sequence clustering algorithm, to resolve a convoluted sample into meaningful subsets and link it to prior knowledge of biological activity. In addition to a simple user-interface (Supplementary Fig. 1A), RoLiM provides multiple ways for users to extract protease and protease substrate patterns from complex datasets, and to visualize the result based on quantitative and statistical assessment.

**Figure 3.**
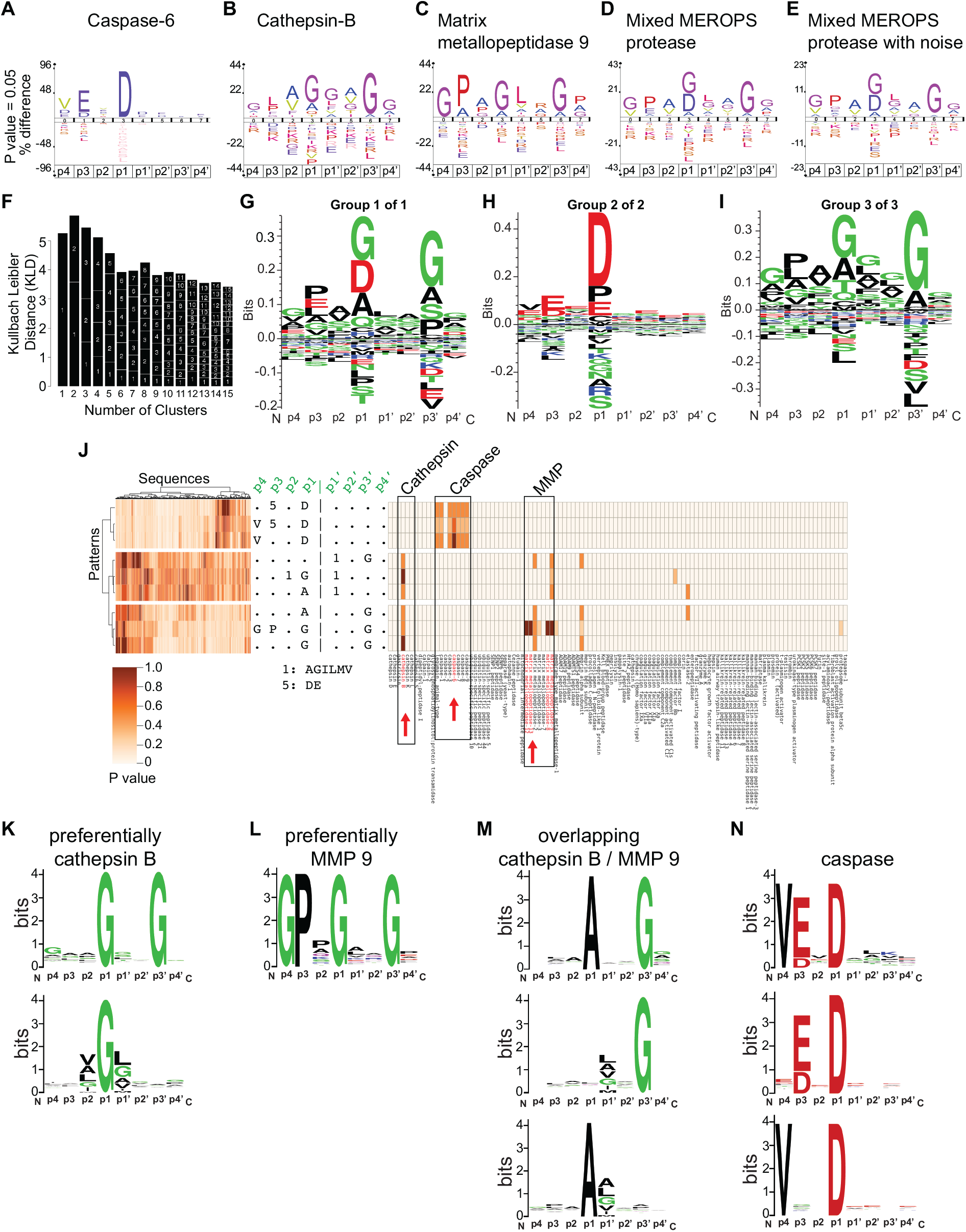
Comparison of GibbsCluster and RoLiM on a mixed MEROPS protease substrate set. **A-E.** Aligned sequence logo for (A) 194 caspase-6 substrates, (B) 469 cathepsin-B substrates, (C) 301 substrates of MMP-9, (D) mix of 964 sequences from A, B and C. (E) sequences in D and an additional 1000 theoretical random sequences to simulate noise. See Methods for more details of the analysis. **F**. Gibbs Cluster analysis of 1951 mixed protease sequences with background noise. GibbsCluster stacked bar chart shows Cluster 2 is the selected solution with the highest average system KLD. **G, H, I.** Logo map of patterns selected for 2 cluster solution shows enriched features consistent with (G) co-clustering of cathepsin B and MMP-9 substrates and (H) caspase-6 sequences respectively. (I) logo map for the third cluster did not resolve cathepsin B and MMP-9 co-clusters. **J.** RoLiM clustermap displaying patterns detected in mixed MEROPS sample containing caspase-6, cathepsin B and MMP-9 substrates. Each column represents a sequence in the data set and each row represents a pattern detected by RoLiM. Both rows and columns are clustered and the matching dendograms are shown to the left and above the clustermap respectively. Pattern distances are encoded as length in the dendograms. Next to the clustermap is a heatmap displaying the patterns detected along with protease specificity annotation from the pre-computed MEROPS protease substrate patterns included in RoLiM. **K-N.** Sequence logos for enriched patterns identified by RoLiM for proteases in a mixed MEROPS protease sample subset.

### Sensitive detection of cancer and treatment specific protease activities in biofluids

Lastly, we re-analyzed two peptidomic and terminomic datasets to demonstrate the application of RoLiM to the identification of proteolytic activities in extracellular fluids of cancer patients. In a multi-centre study on bladder cancer, Frantzi et al^17^ analyzed 1,357 patient urine samples by mass spectrometry-based proteomics and validated urine-based biomarker panels for detecting urothelial bladder cancer. A case-control comparison between 341 primary urothelial bladder cancer cases and 110 urologic controls identified 382 potential biomarkers (reported in Supplementary Table 3 in ^17^). RoLiM analysis on the 382 peptide biomarkers showed a distinct enrichment of sequence patterns surrounding the peptide N and C termini that were in line with matrix metalloproteinase (MMP) specificity motifs (Figs. 4A and 4B). Perhaps due to the absence of a suitable software, the authors did not thoroughly investigate reported peptide sequences for protease substrate motifs. They however report the presence of peptide fragments from a metalloproteinase domain-containing protein 22 (ADAM22), and metalloproteinase with thrombospondin motifs 1 (ADAMTS1). In line with the author’s report, we found evidence for increased MMP cleavage of protein substrates in patients with urothelial bladder cancer compared to disease-free controls. MMP9 and MMP10 were also part of a reported 10-marker ELISA-based assay that enabled discrimination (AUC = 0.85, 95% CI, 0.80-0.91) of urothelial bladder cancer patients from healthy and benign controls^18^ further supporting the assumption that a majority of the 382 peptide biomarkers were a result of cancer associated MMP activity.

**Figure 4.**
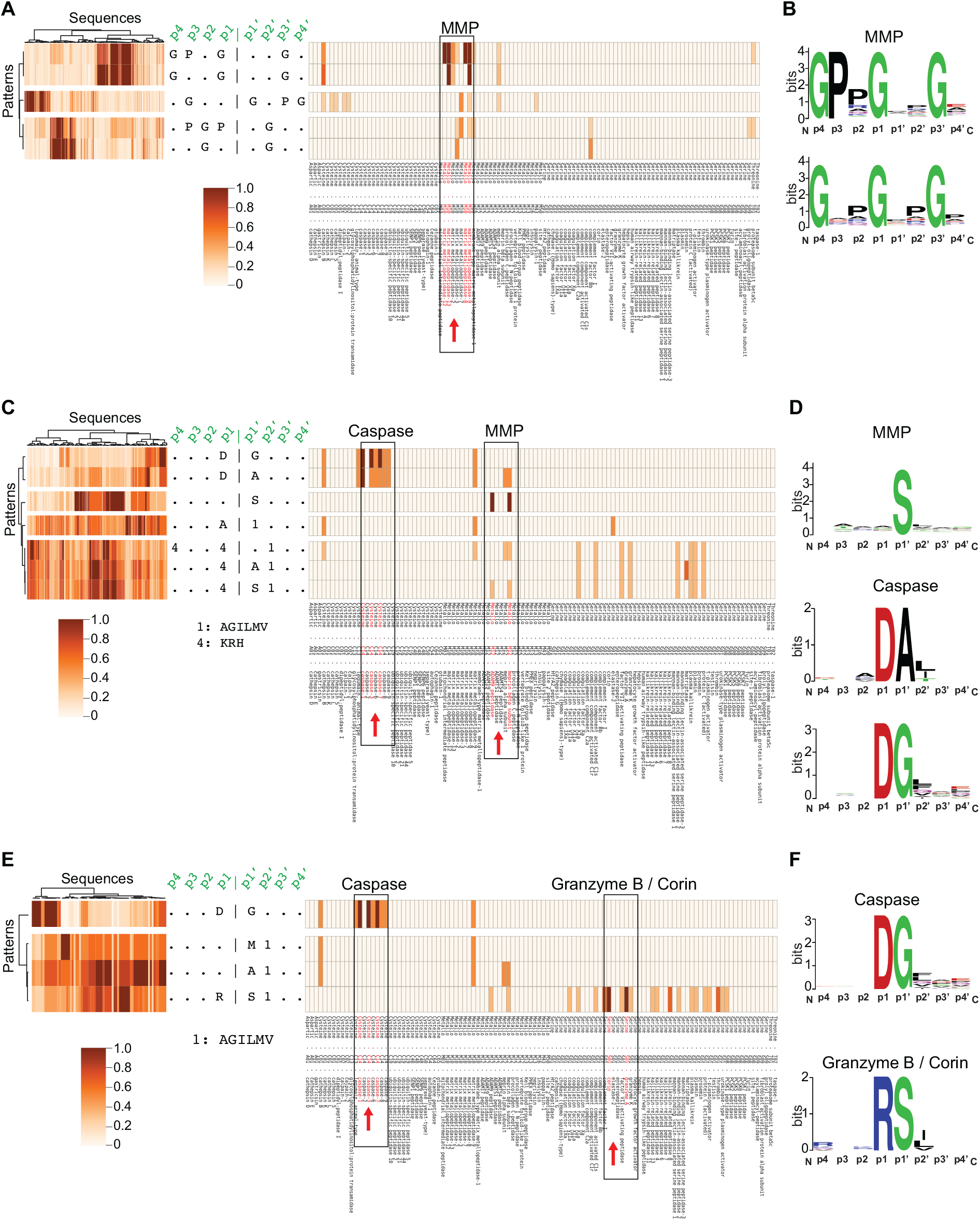
RoLiM enables robust and accurate characterization of protease activities in extracellular fluids from cancer patients. **A**. RoLiM analysis on 382 potential peptide biomarkers for primary urothelial bladder cancer. Clustermap shows patterns enriched in peptide data, and heatmap depicts the main proteases (MMPs) annotated with these patterns. **B**. Sequence logos from aligned peptides with MMP motif. **C, D**. RoLiM analysis on 301 sequences reported by Wiita et al., in all patient cancers (C) and the sequence logos for predicted caspase and MMP proteases (D). **E, F.** RoLiM analysis on 90 sequences uniquely identified by Wiita et al., in only cancer samples (E) and the sequence logos for predicted caspase and granzyme B / Corin proteases (D).

A second study investigated the induction of cellular apoptosis during chemotherapy^19^. The detection of proteolytic fragments released into body fluids by apoptotic tumor cells could offer novel indicators of chemotherapy-induced cell death. Using an N-terminomic approach, Wiita et al.,^19^ found distinct biological signatures of apoptosis in the plasma of hematologic malignancy patients after chemotherapy. These proteolytically generated fragments were notably absent in normal blood. We re-analyzed the non-redundant peptide sequences reported by the authors (Dataset S1 in Witta et al, 2014) for all patient cancers combined (301 sequences), for patient cancer-specific signatures (90 sequences absent in normal controls), and for N termini unique to normal controls. Consistent with the author’s findings, RoLiM clearly identified a caspase D|G and D|A (p1|p1’) signature in a combined analysis of fragments from all cancer patients (Figs. 4C and 4D). RoLiM also revealed a single amino acid (x|S) pattern from MMPs as well as a weak motif for multiple serine proteases including granzymes. Interestingly, this apoptosis related to granzyme signature (R|S[LIV]) was more pronounced in cancer-specific fragments, along with a distinct caspase motif (Figs. 4E and 4F) supporting the sensitivity of the RoLiM algorithm to even detect minor disease related activities among normal proteolytic background activities. The granzyme signature was not described in the paper, and points to the relevant application of RoLiM for unbiased investigation of protease substrate motifs. RoLiM did not detect any overrepresented pattern in the subset of 83 N termini detected in only normal controls.

## Discussion

RoLiM was developed to expand on key functionalities of existing motif discovery and protease prediction tools, and deliver more robust results. The development and characterization of RoLiM is detailed in Smith et al., 2021^9^. Unlike existing tools including Proteasix, Gibbs clustering and Motif-X/Momo, our generalized workflow circumvents the need for a sequence position descriptive input file, or a fixed central residue, and is optimally suited for the analysis of proteolytic data sets. Furthermore, the inclusion of a positional weighting term in the enrichment calculation emphasizes low diversity positions, thus increasing the ability to deconvolute different activities that often have determining sequence features in the same position^9^. Dynamic background frequency adjustment ensures that individual amino acid frequencies are averaged across a background proteome while correcting for the removal of amino acids from the foreground. Vigorous multiple testing is performed where p-values are Bonferroni corrected based on the total number of residues that could be detected at a particular point in the data flow of the algorithm^9^. In addition to enriched individual amino acids, the RoLiM algorithm detects cumulative enrichment of groups of biochemically similar amino acids in the same position, here described as compound residues. This functionality is essential since protease cleavage site specificities are often determined by the general physico-chemical properties of flanking amino acids around the P1 residue rather than a single specific amino acid.

We demonstrated that, in contrast to machine learning based algorithms like Proteasix, RoLiM is not limited to the identification of previously characterized activities. This utility of RoLiM is highlighted in our analysis of time-course data profiling proteolysis in SARS-CoV-2 infected human cell lines. Results from RoLiM analysis revealed more biologically-relevant information that was not limited to pre-existing protease data in MEROPS. While our tool supports the identification of unknown protease substrate patterns in convoluted datasets, additional validation would be needed to verify the protease activity in biological samples.

We also confirmed that RoLiM outperforms alternative unsupervised algorithms, such as Gibbs clustering, that fail to ignore noise and perform poorly in discerning highly similar activities. RoLiM provides null statistics for data lacking overrepresented patterns rather than randomly assigning clusters to likely motifs. Additionally, the descriptive statistics and visual outputs provided by our algorithm supports easy and clear interpretation of individual protease patterns and explicitly conveys the wealth of cleavage event data stored within the MEROPS database. This sets RoLiM apart from other existing motif predictors and sequence clustering software.

Protein fragments, including disease-associated peptides exist in body fluids^3,20,21^. In hematologic malignancies, cellular components in the interstitial space is bathed in a different milieu of substances in comparison to the same extracellular space in normal conditions^22,23^. This is mostly due to the presence of tumor infiltrating cells and the effect of their interactions with other cells in the cancer micro environment. Understanding the protease network in hematologic diseases at different timepoints is relevant for the pathophysiology and discovery of potential markers. Our non-deterministic algorithm strongly supports the unbiased exploration of peptide classifiers, and simplifies the detection of additional leads for follow-up validation of protease-substrate relationships. For example, Wiita et al^19^ did not detect the apoptosis related granzyme signature (R|S[LIV]) in their data on all hematopoietic cancers. An indication of this signature is weakly present in our RoLiM data on all cancers and this signature is further confirmed as significantly overrepresented in the cancer-specific data subset.

In summary, our work for the first time establishes a robust pattern detection algorithm for the identification and annotation of protease activities. RoLiM is designed to meet the needs of researchers for detailed exploration of complex peptidomic and terminomic data to draw biological insights and/or generate testable hypotheses. Our re-analyses of diverse data identify SARS-CoV-2 protease activity in infected cells and link proteolytic products in liquid biopsies to cancer and treatment specific protease activities, demonstrating the profound insights that can be obtained from RoLiM analyses.

## Experimental Section

### Database information

#### Swiss-Prot data

Context sequences were imported from an offline copy of the Swiss-Prot human proteome^24,25^ stored in FASTA format. These sequences were used for the extension of input peptides by mapping peptides to the proteome using a combination of protein accession numbers and regular expression formats. The human Swiss-Prot proteome is also used by default to calculate expected amino acid frequencies. A similar approach will work for other organism-specific Swiss-Prot proteome.

#### MEROPS data

An offline copy of the MEROPS database version 12.0 was used^11^. The proteolytic cleavage event data stored in MEROPS were used to generate data sets of protease substrates. These sequences were retrieved from the database, partitioned by protease, and analysed as input data sets using default algorithm settings. RoLiM was used to detect enriched patterns in each of these protease substrate data sets. Proteases analysed for patterns were restricted to those flagged as “human active,” in the database, and synthetic sequences were excluded from the analyses. MEROPS is an active and expanding database, therefore the offline copy used by this tool will periodically be rebuilt (including computation and storage of protease substrate patterns) when a new version of MEROPS becomes available.

### Data Input and processing

RoLiM has a simple and intuitive user interface(Supplementary Fig. 1). It accepts a number of foreground data set formats as input. These formats include pre- aligned sequences of equal width which can be supplied as a text file (pre-aligned text file). Peptides and accompanying protein IDs may also be supplied in the form of a text file (text file peptide list). For peptide lists, RoLiM automatically extends and aligns the N- and C- terminal cleavage sites of each peptide based on the selected FASTA file. All sequences in the text file input must be of equal length to allow for the analysis of post translational modifications centered around a particular bond. A FASTA file applicable to the context data can be selected as either the default Swiss-Prot Human FASTA file, or other FASTA file uploaded by the user (Supplementary Fig. 1).

The data processing steps are as described in Smith et al.,^9^

### Comparison RoLiM and Proteasix

Quantified N termini reported in Supplementary Table 1 in Meyer et al.,^12^ were analyzed with Proteasix and RoLiM. For analysis in Proteasix, the data was prepared in the required input format, and analyzed with the protease predictor tool using the default settings. The resulting output is summarized in Supplementary Table 5. Proteasix matches a peptide to multiple proteases with identical cleavage patterns. We selected sequence-protease matches with a specificity threshold >= 95. For each protease, the average N termini intensity was determined by calculating the mean intensity for all peptide sequences (Supplementary Table 1) matched to that protease. The resulting data matrix consisted of proteases (rows) and associated sequence averages across all timepoints (columns). A row-wise normalization to the 0 h control was performed on this data. For each row, the average pattern intensity for 0h M, 0h A, 0h B, 0h C was subtracted from the intensity values at timepoints 6 h, 12 h and 24 h respectively. A heatmap with row-wise hierarchical clustering was plotted using the ggplot’s heatmap.2 function in R.

With RoLiM, N termini were analyzed using the default setting with the following additional changes: p value threshold of 0.05, minimum occurrence set to 5% of the input (4, 7, 8, 17 sequences respectively), width at 8, and centered sequences activated. Overrepresented patterns identified by RoLiM and grouped according to each timepoint are listed in Supplementary Table 6. Each pattern was matched to the corresponding sequences in Supplementary Table 1 based on its P4 – P4’ alignment. The resulting matrix contained a list of all quantified N termini from Meyer et al, each with an associated RoLiM pattern. The abundance of a pattern across all timepoints was determined using the average intensity of all peptides linked to that pattern. This generated a matrix containing all identified patterns (rows), and the mean intensity of matched peptides per pattern across all timepoints (columns). A row-wise normalization to the 0 h control was performed on this matrix. For each row, the average pattern intensity for 0h M, 0h A, 0h B, 0h C was subtracted from the intensity values at timepoints 6 h, 12 h and 24 h respectively. The resulting data with normalized ratios to timepoint 0 was used as input for hierarchical clustering plots in R using ggplot’s heatmap.2 function.

### Comparison of RoLiM and Gibbs Clustering

An analysis of the frequency of false positive detection by our algorithm was already conducted and is described in Smith et al.,^9^.

#### Generation of *in silico* datasets

An *in silico* set of 1000 sequences containing non-fixed positional amino acids was randomly constructed from the Swiss Prot human proteome using NumPy arrays^26^ in Python. This data (Supplementary Table 3) was supplied as input, and Gibbs clustering was run over a range of 1 to 15 possible clusters respectively (the minimum and maximum settings for the web interface). A motif length of 8 (the full length of the input sequences) was selected, the simple shift move of the algorithm which attempts realignment of the dataset was disabled, and the trash cluster for outlier rejection was enabled at 0. All other settings were kept at their default values. The same data was supplied as input for RoLiM using the default settings, a width of 8 and centered sequences checked. The Human Swiss Prot Fasta was supplied as context proteome.

#### Mixed protease data set

This data set was generated using a combination of human protease substrate sequences from the MEROPS database. This included 194 caspase-6 substrates, 469 cathepsin B substrates, 301 matrix metalloproteinase 9 substrates, and 987 random Swiss Prot human protein sequences (Supplementary Table 4). Gibbs clustering was run over 1 to 15 possible clusters, motif length of 8 (the full length of the input sequences) was selected, the simple shift move of the algorithm which attempts realignment of the dataset was disabled, and the trash cluster for outlier rejection was enabled at 0. All other settings were kept at their default values. The same data was supplied as input for RoLiM using the default settings, a width of 8 and centered sequences checked. The Human Swiss Prot Fasta was supplied as background proteome.

### Urinary peptidome data

We analyzed N and C terminally extended, non-redundant sequences reported by Frantzi and colleagues (Supplementary Table 3 in ^17^). RoLiM default settings of p value < 0.001, minimum frequency of 20 was used. A width of 8 was maintained and centered sequences option was checked.

### Hematologic cancer data

Non-redundant sequences reported for all cancers combined, or for cancer-specific patients in Wiita et al (Supplementary Table S1 in ^19^) were analyzed. RoLiM settings were p value cut-off of 0.001, minimum frequency of 5., width at 8, and centered sequences activated.

### Data output files

Logo maps used were generated in WebLogo^27^. RoLiM automatically generated logo maps for data analyzed with the algorithm. Logomaps were used to visualize the amino acid distribution of each subset of aligned input sequences exactly matching one of the detected patterns.

### Supporting Information

Additional details for experimental analyses are provided and include input data, figures and results.

## Abbreviations

RoLiM: RObust LInear Motif Deconvolution
PTM: Post-translational modification

## Acknowledgements

We thank Lorenz Nierves for discussions and constructive feedback, and Enes K. Ergin for contributions with database literature search.

## Competing Interests

All authors declare no competing interests

## Funding sources

This work was partially supported by grants from the Natural Sciences and Engineering Research Council of Canada (NSERC) (#RGPIN-2018-05645), the Michael Cuccione Foundation and the BC Children’s Hospital Foundation (to P.F.L.). A.C.U. was supported by fellowships from the BC Children’s Hospital Research Institute and Michael Cuccione Childhood Cancer Research Program and T.G.S was supported by a NSERC CREATE studentship. P.F.L was supported by the Canada Research Chairs program, the Michael Smith Foundation for Health Research Scholar program and the Michael Cuccione Foundation.

## Notes

### Competing Interest Statement

The authors have declared no competing interest.

