## Supplementary Information for "Sensitive identification of known and unknown protease activities by unsupervised linear motif deconvolution"

#### Supporting Information

##### Supplementary Figures:

Supplementary Figure S1. Analysis of N terminome of SARS-Cov2 infected human cell line.
Supplementary Figure S2. MEROPS annotation function in RoLiM.
Supplementary Figure S3. GibbsCluster analysis on a set of 1000 random Swiss-Prot sequences.
Supplementary Figure S4. Logo Maps for Gibbs Cluster solution on 1000 random Swiss-Prot sequences

##### Supplementary Tables:

Supplementary Table S1. Quantified N termini reported in Meyer et al 2020 Table S1

Supplementary Table S2. MEROPS human proteases, the number of validated physiological substrates per protease, and count of literature citations and experimental reports for each protease

Supplementary Table S3. 1000 in silico random sequences from the Swiss Prot human proteome analyzed in GibbsCluster and RoLiM

Supplementary Table S4. Mixed MEROPS protease data set. Input sequence for mixed data set plus noise, and individual protease

Supplementary Table S5. Summary of result from Proteasix analysis of Meyer et al S1 data

Supplementary Table S6. List of overrepresented ROLiM patterns detected for N termini enriched across virus infection time points, as reported in Meyer et al Table S1.

#### Supplementary Figures

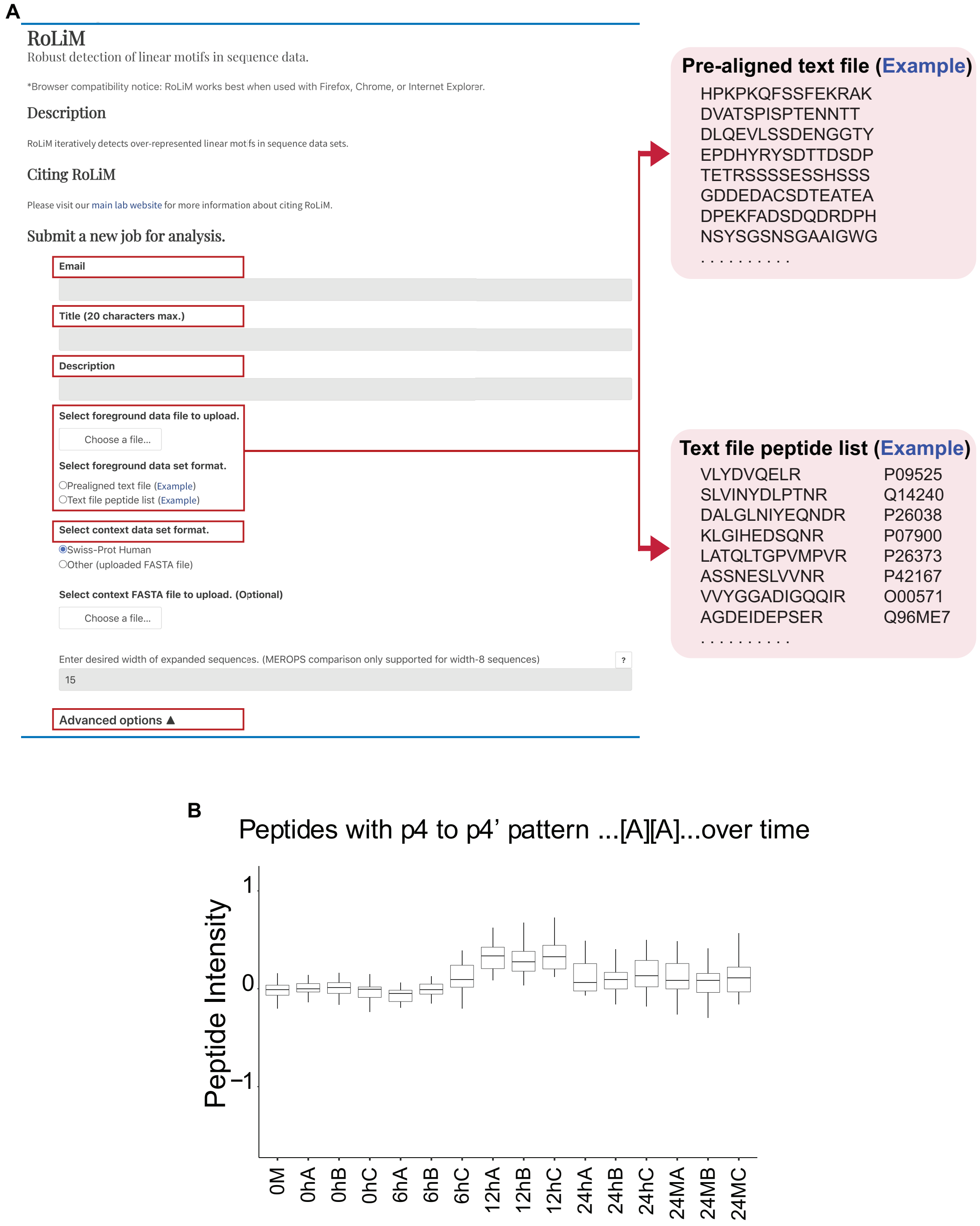


Supplementary Figure S1. Analysis of N terminome of SARS-Cov2 infected human cell line.

**A.** Intuitive RoLiM user interface.

**B.** Bar plot showing the normalized intensity of peptides with the ...[A][A]... pattern in infected and non-infected cells at all timepoints studied.

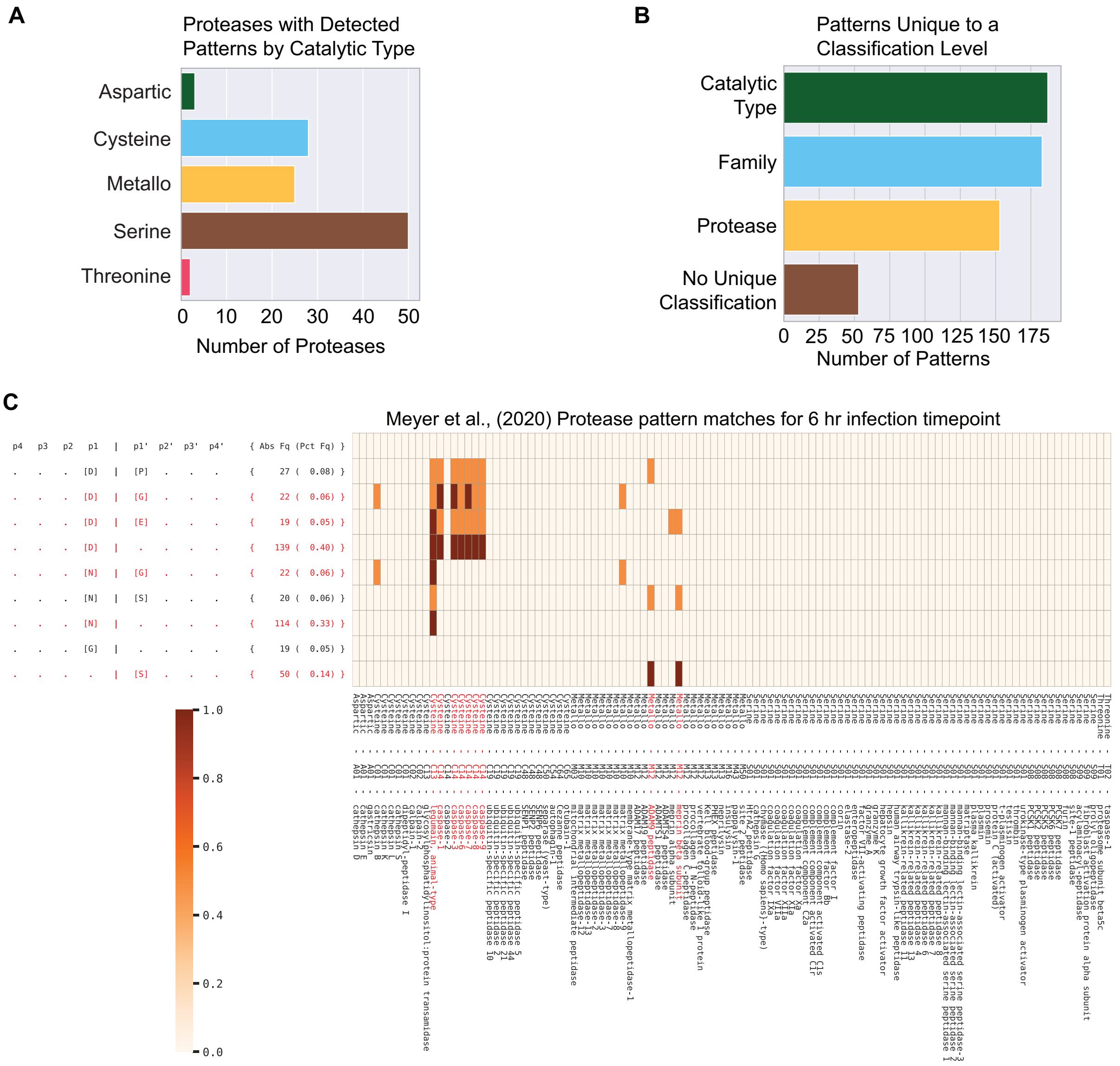


### Supplementary Figure 2. MEROPS annotation function in RoLiM.

**A.** Characterization of MEROPS protease substrate data using RoLiM. The 261 patterns detected by RoLiM are grouped according to catalytic type.

**B.** Classification of RoLiM patterns. Unique patterns classified based on catalytic type, protease family, individual protease, or with no defined classification level in MEROPS.

**C.** RoLiM protease pattern output. An example of a protease pattern heatmap showing enriched sequences mapped to MEROPS curated protease patterns.

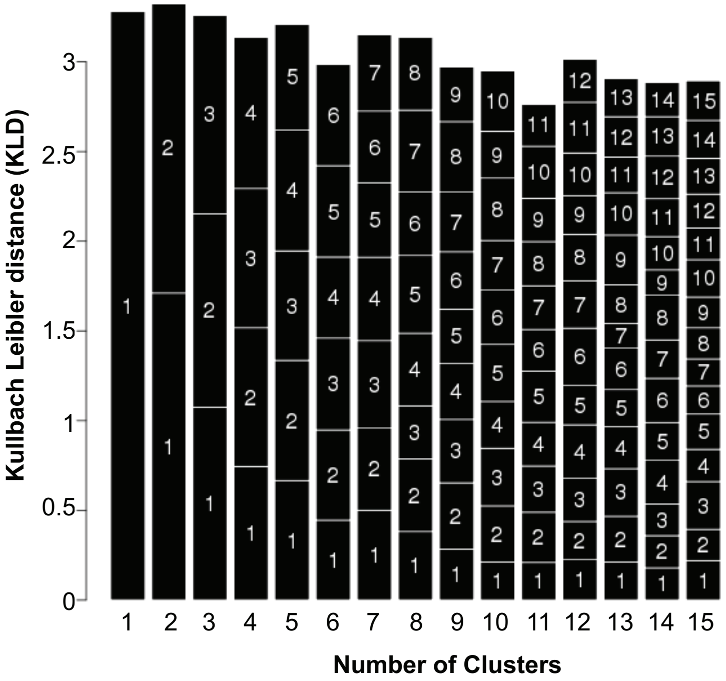


Supplementary Figure 3. GibbsCluster analysis on a set of 1000 random Swiss-Prot sequences.

Stacked bar chart showing the mean KLD for each cluster in a set of solutions ranging from 1 cluster to 15 clusters. Gibbs Cluster selects the solution with the highest average system KLD (cluster 2). The algorithm selected a solution of 2 clusters with an final average KLD of 2.893. Both logo maps for this solution are also represented.

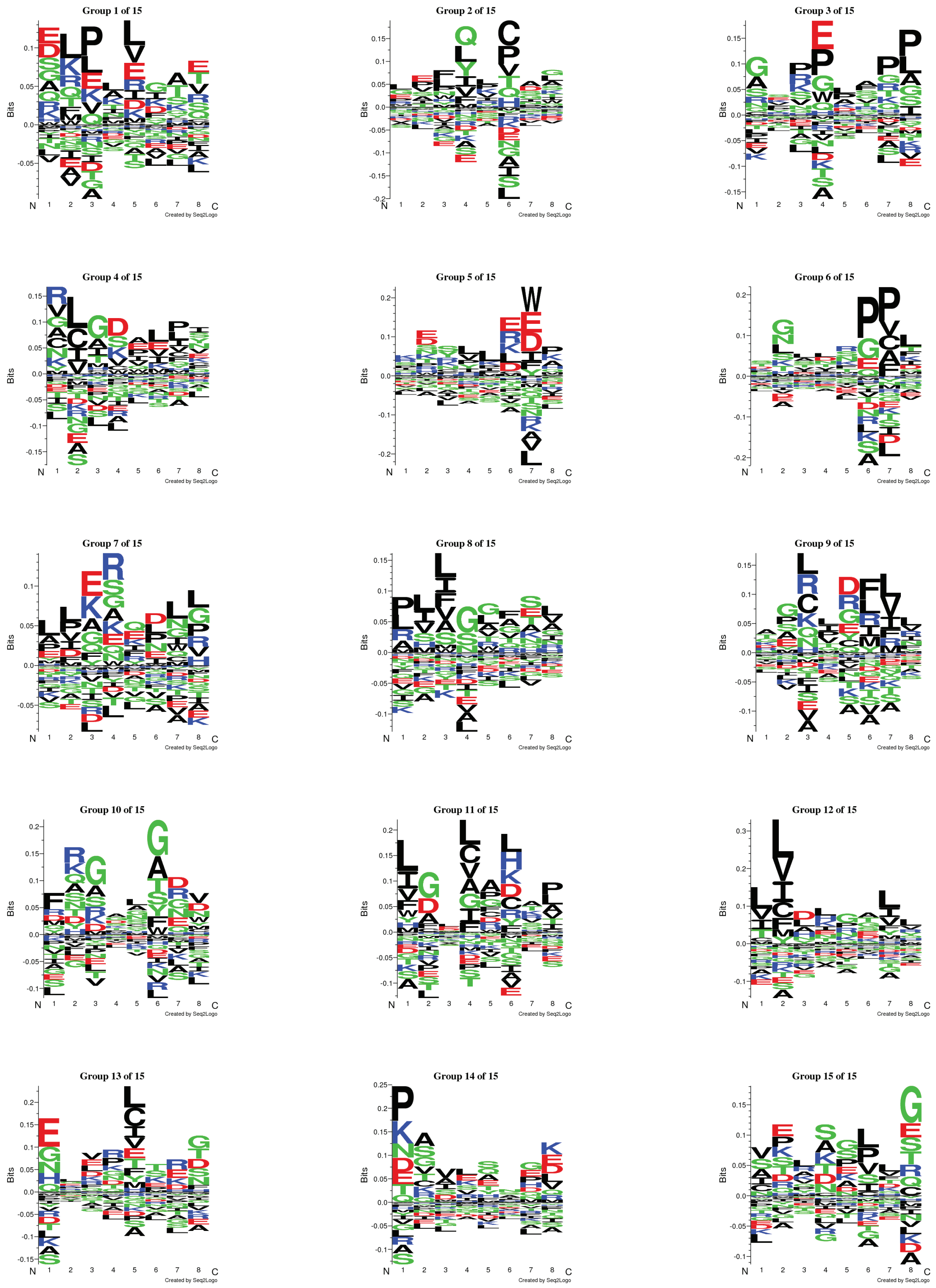


Supplementary Figure 4. Logo Maps for Gibbs Cluster solution on 1000 random Swiss-Prot sequences. The algorithm selected a solution of 2 clusters from a range of 1 to 15 clusters. Sequence Logo Maps show the aligned sequence patterns for each cluster.

**Supplementary Tables**

Supplementary Table S1. Quantified N termini reported in Meyer et al 2020 Table S1

Supplementary Table S2. MEROPS human proteases, the number of validated physiological substrates per protease, and count of literature citations and experimental reports for each protease

Supplementary Table S3. 1000 in silico random sequences from the Swiss Prot human proteome analyzed in GibbsCluster and RoLiM

Supplementary Table S4. Mixed MEROPS protease data set. Input sequence for mixed data set plus noise, and individual protease

Supplementary Table S5. Summary of result from Proteasix analysis of Meyer et al S1 data

Supplementary Table S6. List of overrepresented ROLiM patterns detected for N termini enriched across virus infection time points, as reported in Meyer et al Table S1.
